# GDF15 mediates renal cell plasticity in response to potassium depletion

**DOI:** 10.1101/2022.12.27.521983

**Authors:** Samia Lasaad, Christine Walter, Chloé Rafael, Luciana Morla, Alain Doucet, Nicolas Picard, Anne Blanchard, Yves Fromes, Béatrice Matot, Gilles Crambert, Lydie Cheval

## Abstract

A low potassium (K^+^) intake is a common situation in the population of the Westernized countries where processed food is prevalent in the diet. Here, we show that expression of GDF15, a TGFβ-related growth factor, is increased in renal tubular segments and gut parts of mice in response to low-K^+^ diet leading to a systemic elevation of its plasma and urine concentration. In human, under mild dietary K^+^ restriction, we observed that urine GDF15 excretion is correlated with plasma K^+^ level. Conversely to WT mice, adaptation to K^+^ restriction of GDF15-KO mice is not optimal, they do not increase their number of type A intercalated cell, responsible for K^+^ retention, and have a delayed renal K^+^ retention, leading to early development of hypokalemia. This renal effect of GDF15 depends on ErBb2 receptor, whose expression is increased in the kidney collecting ducts. We also observe that, in the absence of GDF15, the release of K^+^ by the muscles is blunted which is compensated by a loss of muscle mass. Thus, in this study, we showed that GDF15 plays a central role in the response to K^+^ restriction by orchestrating the modification of the cell composition of the collecting duct.

## Introduction

The modern Western diet is characterized by consumption of processed food with high intakes of proteins, high-fat dairy products and high-sugar drinks correlating with development of metabolic disorders (diabetes, obesity). This type of diet has also strong but often underappreciated consequences on the electrolyte balances. Indeed, modern human eating typical Western diet produces approximately 50 mmol of acid/day whereas our ancestors were considered to be net base producers (1). In parallel, the consumption of salt (NaCl) and potassium (K^+^) have been completely modified, switching from a rich-K^+^/low NaCl diet in the hunter-gatherer population to the opposite in the modern, westernized population (2). In a recent study, we showed that the median daily Na^+^ and K^+^ intakes in young Parisian males are 136 mmol and 57 mmol/day, respectively (3) which are far from the recommended values by the dietary reference intakes (maximal Na^+^ intake 65 mmol/day and minimal K^+^ intake 120 mmol/day). Fortunately, we have the ability to cope with these changes that occurred very recently in our eating habits thanks to our kidneys. However, these long-term dietary modifications may also contribute to the development of diseases such as hypertension (4) and are correlated with higher risk of mortality (5) when they encounter favorable genetic background with polymorphisms that would have remained silent under more “natural” diet. The lack of K^+^ in the diet is a situation well-known to induce a global adaptation involving coordinated regulatory processes (6). The ability of muscle to release K^+^ (internal balance) is of particular importance for rapidly modulating the plasma K^+^ concentration (7). In parallel, the kidney decreases its ability to secrete K^+^ (through the inhibition of K^+^ channel ROMK, for review see (8)) and activates K^+^ retention pathways through a progesterone-dependent stimulation of the H,K-ATPase type 2 (9) and the increase of the number of A-type intercalated cells (AIC) in collecting duct (10, 11).

The cellular plasticity of the collecting duct is a fascinating process allowing the kidney to modify its structure to respond to ionic stress through mechanisms involving either transdifferentiation from another cell type of the collecting ducts and/or direct cell proliferation. Our group has identified that the growth differentiation factor 15 (GDF15) triggers the proliferation of AIC in response to acidosis (12, 13). GDF15 is a peptide belonging to the TGFβ family which has been reported to have multiple physiological effects (14). Regarding the renal production of GDF15, in addition to the collecting duct, as mentioned above, it has also been established that it is produced by a subset of proximal tubule cells in mice mimicking Cockayne syndrome (15). It is now strongly suggested that GDF15 is a sentinel mediator (16) that responds to stress situation involving mitochondrial defects (17–19) and is found increased in the circulation and urine following many pathological states, mainly in cancer. GDF15 is also involved in the control of energy metabolism, being overexpressed in obese condition (20, 21) and having anorectic properties (22, 23). GDF15 is also found increased in aging (24) where it could protect against inflammation and tissue injury but also to muscle wasting at this stage. Interestingly, GDF15 has also been identified as one of the most upregulated genes in kidney collecting ducts in response to dietary K^+^-restriction (10) but its role in this particular situation was yet unknown. In this study, we show that GDF15 plays a central role in the control of the K^+^ balance by adapting the cell composition of the collecting duct and found that the inability to increase AIC, as observed in GDF15-KO mice, leads to loss of muscle volume.

## Results

### GDF15 level is increased in response to dietary K^+^-restriction

We showed that urine GDF15 is increased (Figure 1A) by K^+^ restriction following a kinetic that ends to a plateau at day 3-4. At this plateau, the level of GDF15, normalized to the creatinine, is 3-time higher than in control diet (877±104 ng/mmol at day 4 vs 310±25 ng/mmol in control diet). In parallel, the level of GDF15 in the plasma (Figure 1B) is also 2-time higher in mice under low-K^+^ diet for 4 days than in control diet (140±14 vs 69±2 pg/ml, respectively) indicating that the stimulation of GDF15 is more systemic than previously envisaged. As shown in the Figure 1C, the expression of GDF15 is increased to a different extent in all nephronic segments from mice under low-K^+^ diet. Since the stimulation of GDF15 is more systemic than previously envisaged, we therefore extended our investigation to muscle tissues and intestine which are known to be involved in K^+^ metabolism. As shown in Figure 1D, we measured the tissue content of GDF15 by ELISA analysis and we showed that it is significantly increased in the ileum and the colon of mice under K^+^ restriction but not in muscle.

**Figure 1:**
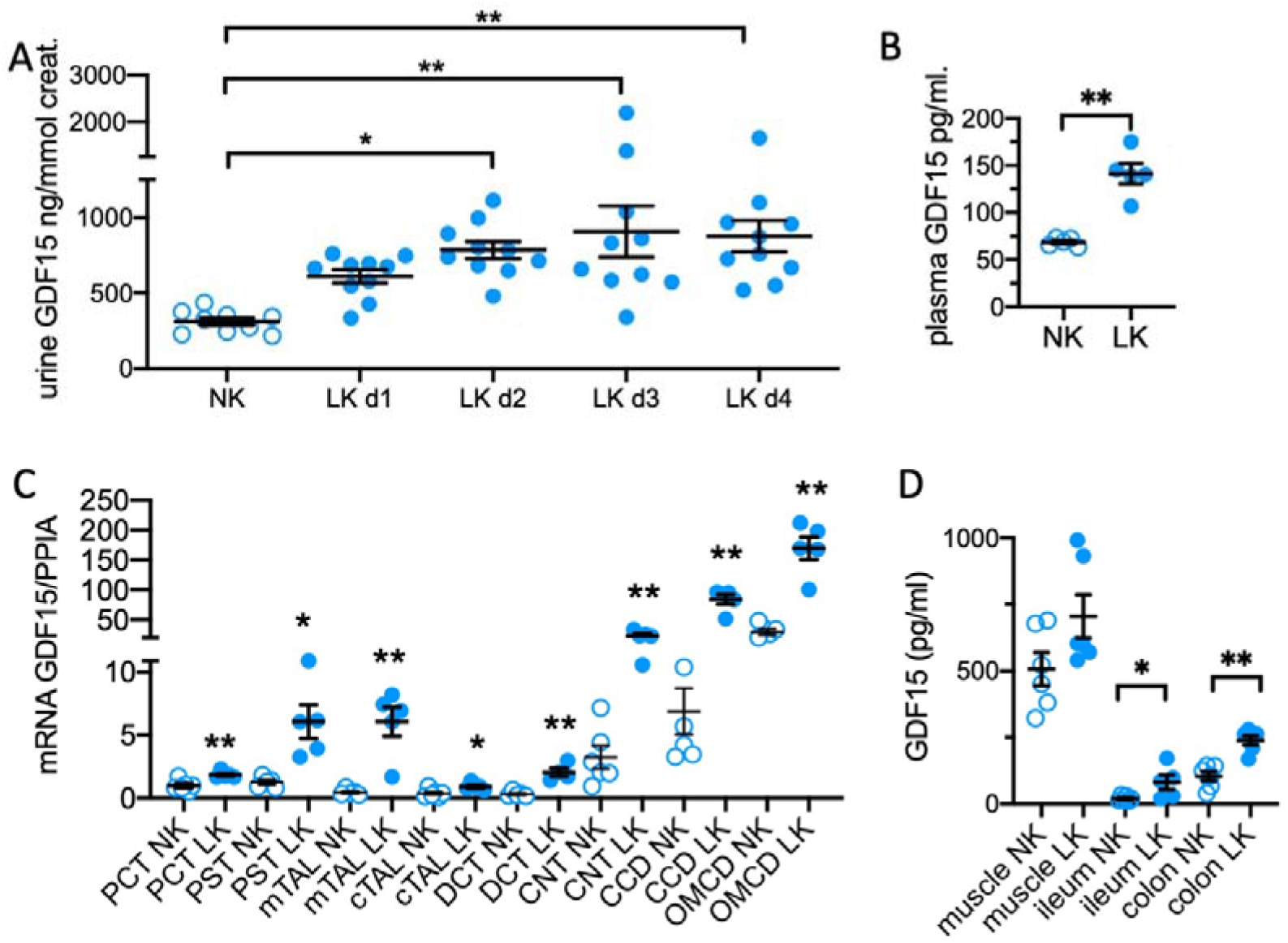
GDF15 is produced in response to K^+^ restriction. **A/** Urine GDF15 normalized by urine creatinine under control condition (NK, empty symbols) and at day 1 to 4 of a low-K^+^ diet period (LK, filled symbols). Results are shown as mean±SEM (n=10) and analyzed by a one-way ANOVA test (p<0.01) followed by a Dunett’s multiple comparison test with NK group as a control group (** p<0.01; * p<0.05). **B/** Plasma GDF15 level in mice under normal diet (NK, empty symbols) or K^+^-restriction for 4 days (LK, filled symbols). Results are shown as mean±SEM (n=4-5) and analyzed by an unpaired Student t test (** p<0.01). **C/** mRNA expression of GDF15 in isolated proximal convoluted tubules (PCT), proximal straight tubules (PST), medullary and cortical thick ascending limb (m and cTAL), distal convoluted tubule (DCT), connecting tubule (CNT) and cortical or outer medullary collecting duct (CCD and OMCD) of mice under normal diet (NK, empty symbols) or K^+^-restriction for 4 days (LK, filled symbols). Results are shown as mean±SEM (n=5) and analyzed by comparing the effect of the diet on each segment independently of the others by a Mann-Whitney test (** p<0.01; * p<0.05). **D/** GDF15 protein expression was measured in muscle (gastronecmius), ileum and distal colon tissues by ELISA in mice under normal diet (NK, empty symbols) or K^+^-restriction for 4 days (LK, filled symbols). Results are shown as mean±SEM (n=6) and analyzed by a Mann-Whitney test (** p<0.01).

To investigate whether this link between GDF15 and dietary K^+^ restriction is also present in human, we used urine samples obtained in a former study where healthy young men were K^+^-depleted for one week (3). As shown in Figure 2A, the dietary intervention slightly but significantly decreased the plasma K^+^ level from 3.7±0.1 mM to 3.35±0.09 mM (p=0.039). When we analysed and compared the urine excretion of GDF15 in these subjects under normal diet or after a week of K^+^-depletion, we did not observe any significant difference (Figure 2B). Taking a closer look at the relationship between the presence of GDF15 in urine and the plasma K^+^ level of the different subjects, we found that under normal K^+^ diet, there was no correlation (Figure 2C). However, after dietary K^+^ restriction the urine excretion of GDF15 is significantly correlated to the plasma K^+^ level (Figure 2D).

**Figure 2:**
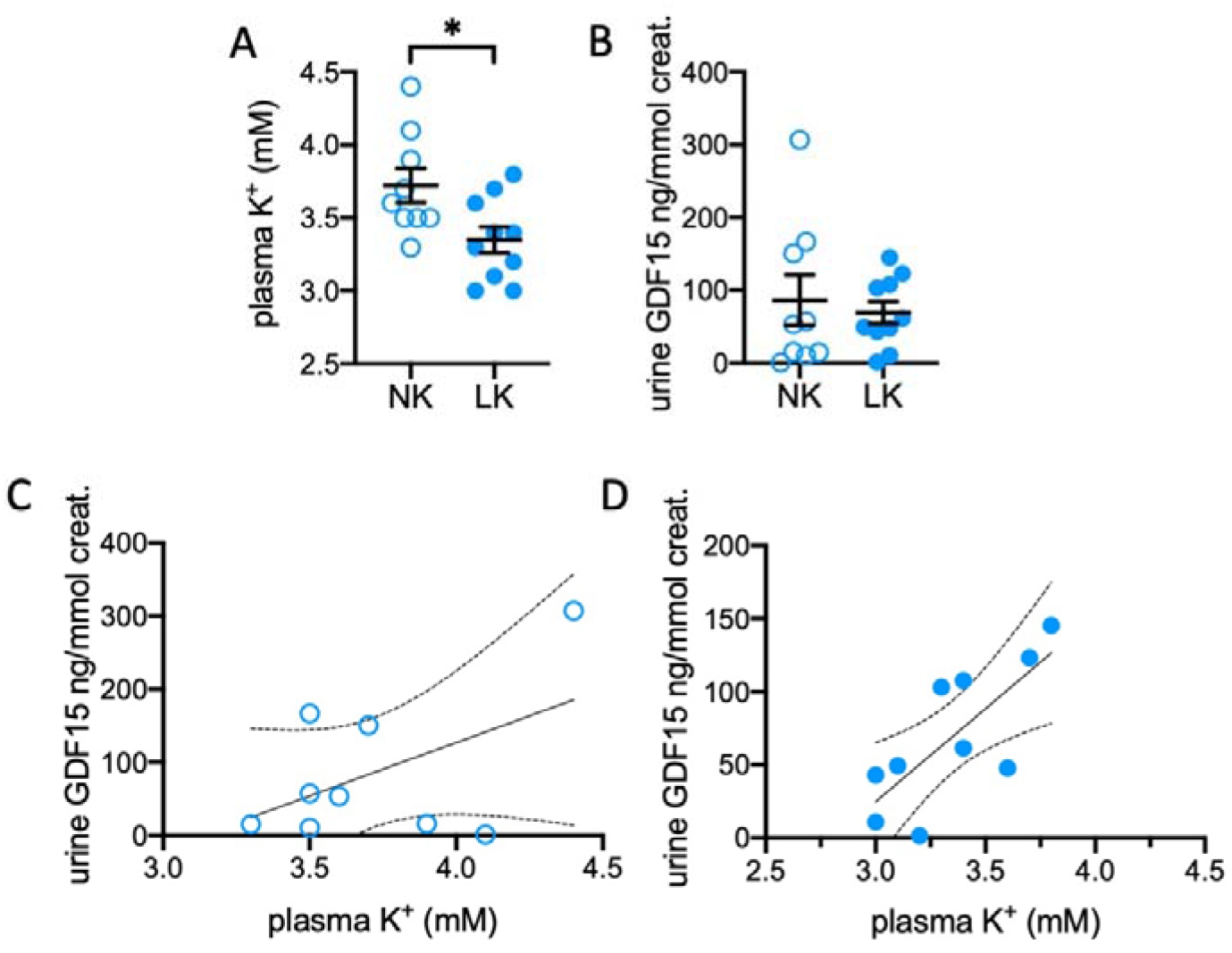
Urine GDF15 level correlates with plasma K^+^ value in human under dietary K^+^ restriction. The groups of healthy volunteers under normal diet (NK, empty symbols) or K^+^-restriction (LK, filled symbols) were already characterized in (3). **A/** plasma K^+^ values in NK (n=9) or LK (n=10). Results are shown as mean±SEM and analyzed by a Wilcoxon test (* p<0.05). **B/** urine GDF15 excretion measured by ELISA. Results are shown as mean±SEM and analyzed by a Wilcoxon test (* p<0.05). Correlation between both parameters for NK group (**C**) and LK group (**D**) assessed with a simple linear regression test indicating that the slope is not different from 0 in the NK group with a Pearson R squared of 0.497 whereas it is in LK group (p=0.013) with a Pearson R squared of 0.749.

### GDF15 is involved in the increase of AIC observed in response to LK diet

To quantify the increase of AIC number, we used a simple method consisting in isolation of medullary collecting ducts and labelling of the anion exchanger 1 (AE1), a specific marker of the AIC (25) (Figure 3A). As shown in Figure 3B, the number of cells/mm is neither modified by the K^+^-restriction period nor by the absence of GDF15 (GDF15-KO). However, the percent of AE1+ cells increased from 20.8±0.7% in control diet to 23.7±0.6% (p<0.05) in WT mice (Figure 3C). In GDF15-KO mice, the percent of AIC under control condition is similar to that in WT mice and did not increase in LK diet condition (20.9±0.4% vs 21.2±0.8%, respectively). A characteristic of AIC is to be completely isolated in the tubular epithelium, the observation of two adjacent cells (doublet) is therefore the trace of a recent cell division. As shown in Figure 3D, the number of AIC doublets was significantly increased by a low-K^+^ diet in WT mice (13.8±1.8 doublets/mm and 21.1±1.4 doublets/mm, respectively, p<0.05). In GDF15-KO mice, the number of doublets is similar than in WT mice under control condition and did not increase after 4 days of LK diet. The expression of Cyclin D1 a gene involved in cell proliferation was then followed in isolated OMCD of WT and GDF15-KO mice under either control diet or 2 and 4 days of LK diet. As shown in Figure 3E, cyclin D1 expression in OMCD is similar in both mouse genotypes but is significantly increased after 2 days of LK diet in WT (p<0.01) but not in GDF15-KO mice indicating a delay in cell division rates in the absence of GDF15. Four days after LK, both genotypes expressed the same amount of cyclin D1 in their OMCD.

**Figure 3:**
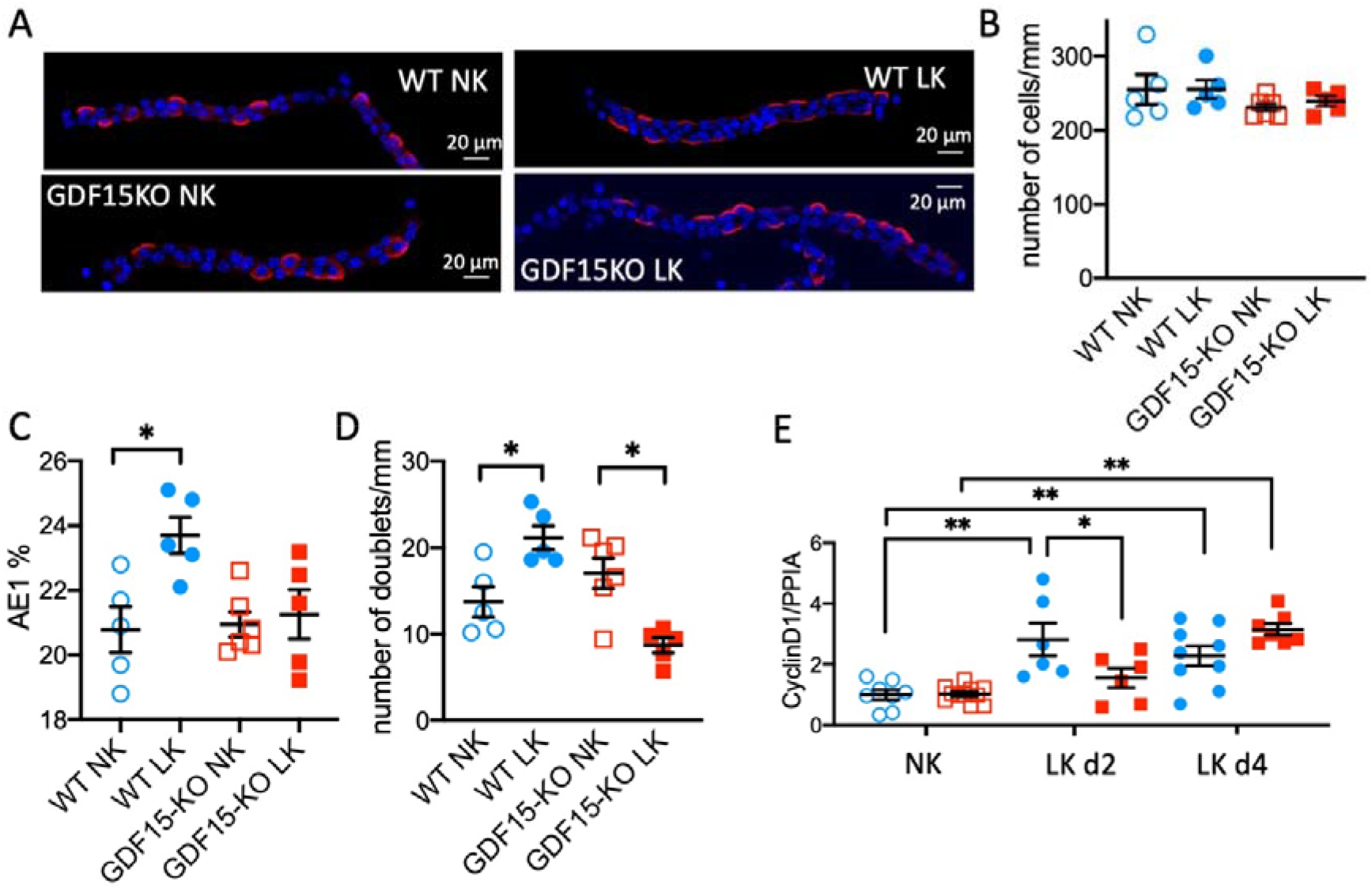
GDF15 is involved in proliferation of ICA under low-K^+^ diet. **A/** Examples of immunofluorescence pictures obtained by confocal microscopy from isolated OMCD from WT (blue circles) or GDF15-KO (red squares) mice under normal (empty symbols) or K^+^-restriction for 4 days (filled symbols) labelled with DAPI (blue) and anti-AE1 antibody (red). The stacks of these images were then used to reconstruct tubules in 3D in order to accurately count the number of nucleus (**B**), the number of AE1+ cells (**C**) and the number of AE1+ doublet (at least two AE1 + cells with cell-cell contact, (**D**). Each symbol represents the mean value of 7-11 reconstructed tubules of the same animal. Results are shown as mean±SEM (n=5-6) and analyzed by a Mann-Whitney test (** p<0.01). **E**/ Cyclin D1 mRNA expression in WT (circles) or GDF15-KO mice (squares) under normal (empty symbols) or K^+^-restriction (filled symbols) for 2 and 4 days (n=6-10). Results are shown as mean±SEM and analyzed by a two-way ANOVA followed by a Sidak’s multiple comparison test. Cyclin D1 expression is strongly affected by the period of the K^+^-restriction (p<0.01) and by the combination of both treatment period and genotype (p<0.01) but not by the genotype itself (p=0.25).

### GDF15 participates in the renal adaptation to K^+^-restriction through an ErbB2-dependent mechanism

To appreciate the involvement of GDF15 in the renal adaptation to a low-K^+^ diet and the physiological consequences of the lack of AIC proliferation, we investigated WT and GDF15-KO mice under K^+^ restriction. The physiological parameters of WT and GDF15-KO mice were reported in Table 1 and showed that prolonged LK diet induced a decrease of weight and food intake in WT that is less pronounced in GDF15-KO mice. Urine volume increased in both genotypes. Measurement of the urine K^+^ excretion of WT and GDF15-KO mice in response to a LK diet (Figure 4A) showed that the absence of GDF15 induced a delayed in the reduction of K^+^-excretion. Thus, at day 1, 2 and 3 of the K^+^-restriction period, the GDF15-KO mice lose between 30 to 100% more K^+^ than WT mice. As shown in Figure 4A *inset*, this delayed renal response has a consequence on the plasma K^+^ level at day 4 of the LK diet, with GDF15-KO mice becoming hypokalemic (3.3±0.1 mM) compared with the control mice (4.3±0.2 mM, p<0.01). On the contrary, WT mice remained with a normal plasma K^+^ value (3.9±0.1 mM) not different from that measured in control diet (4.1±0.1mM). The classical receptor for GDF15 is GFRAL that mediates most of its central nervous effect and metabolic effects (for review, see (26)). However, this receptor is not present in the kidney (25) but we have recently identified that ErbB2 receptor, another receptor of GDF15 was present in the kidney (25). In Figure 4B, we measured ErbB2 expression in whole kidney and isolated segments of the distal nephron from the connecting tubules (CNT) to the cortical (CCD) and medullary collecting duct (OMCD). We observed that ErbB2 is not modified by a LK diet at the level of whole kidney analysis but is significantly increased in the distal nephron by 50-100% in response to K^+^-restriction. This indicates that ErbB2 may be involved in the GDF15-dependent renal response to K^+^-restriction. We tested that hypothesis by treating WT mice under control or LK diet for four days with an antagonist of ErbB2, the mubritinib. As shown in Figure 4C, mubritinib treatment did not modify the number of cell/mm but impeded the LK diet-dependent increase of AIC (Figure 4D-E). Interestingly, the absence of ICA proliferation under a low-K^+^ diet impedes a correct adaptation since we showed that if mubritinib alone did not impact the plasma K^+^ level of mice under control diet, it reduced the plasma K^+^ values of WT mice under LK diet (non-treated mice 4.0±0.2mM vs treated mice 3.3±0.1 mM, p<0.05, Figure 4F). The regulation of cell plasticity in collecting ducts by a GDF15/ErbB2 axis could therefore be an important process to adapt to a dietary K^+^ restriction.

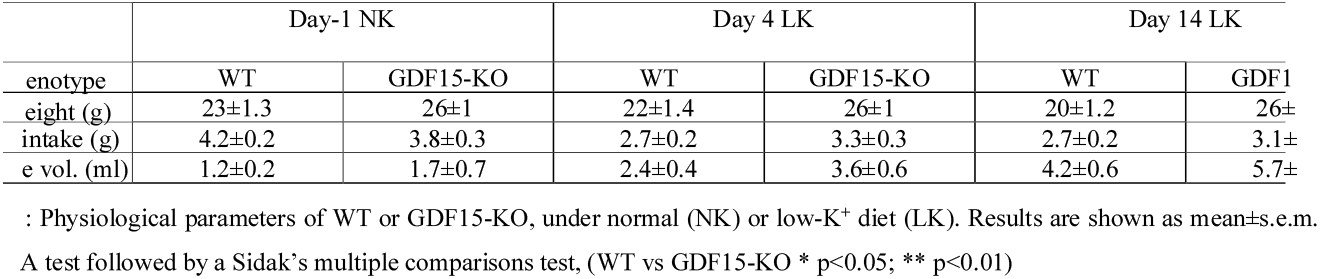

**Figure 4:**
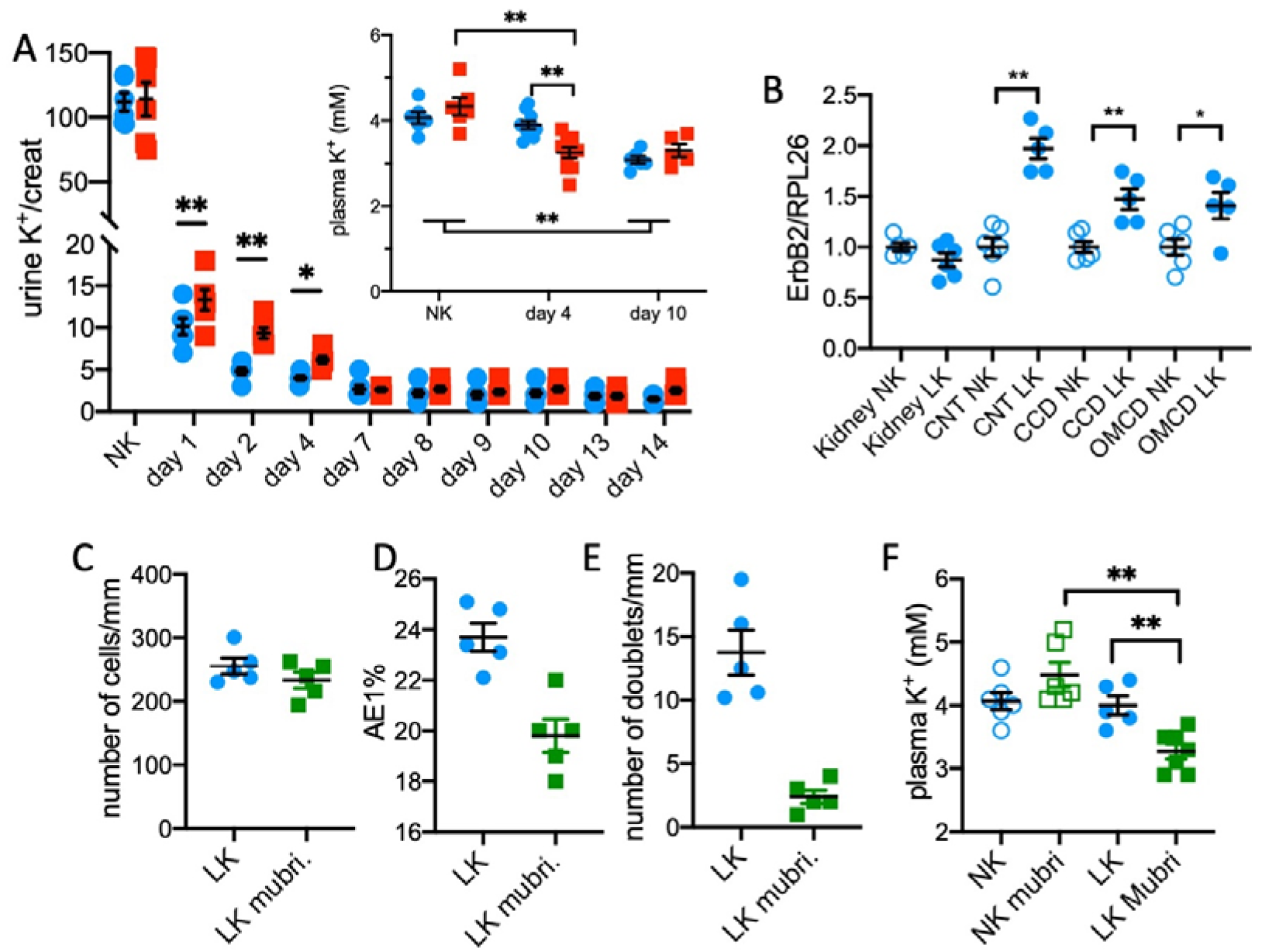
GDF15 regulates K^+^ balance under low-K^+^ diet. **A/** Urine K^+^ excretion normalized by urine creatinine under control condition (NK) and at day 1 to 14 of a low-K^+^ diet (LK) in WT (blue circles) or GDF15-KO (red squares) mice. Results are shown as mean±SEM (n=6) and analyzed by a two-way ANOVA followed by a Sidak’s multiple comparison test (** p<0.01, * p<0.05). ***Inset*** Plasma K^+^ level under control condition (NK) and at day 4 and 10 of a low-K^+^ diet (LK) in WT (circles) or GDF15-KO (squares) mice. Results are shown as mean±SEM (n=6-8) and analyzed by a two-way ANOVA followed by a Sidak’s multiple comparison test (** p<0.01, * p<0.05). **B/** mRNA expression of ErbB2 in total kidney or CNT, CCD and OMCD renal segments of mice under normal diet (NK, empty symbols) or K^+^-restriction for 4 days (LK, filled symbols). Results are shown as mean±SEM (n=5-6) and analyzed by comparing the effect of the diet on each segment independently of the others by a Mann-Whitney test (** p<0.01; * p<0.05). Number of nucleus (**C**), number of AE1+ cells (**D**) and the number of AE1+ doublets (**E**) in WT mice under low-K^+^ diet for 4 days (LK) in the absence (blue filled circles) or the presence (green filled squares) of mubritinib treatment. Each symbol represents the mean value of 7-11 reconstructed tubules of the same animal. Results are shown as mean±SEM (n=4-5) and analyzed by a Mann-Whitney test (** p<0.01). **F/** Plasma K^+^ level in WT mice under control condition (NK, empty symbols) and at day 4 of a low-K^+^ diet (LK, filled symbols) in the absence (blue symbols) or presence (green symbols) of mubritinib treatment. Results are shown as mean±SEM (n=5-7) and analyzed by a two-way ANOVA followed by a Sidak’s multiple comparison test (** p<0.01, * p<0.05).

### The absence of GDF15 induces modification of muscle structure in response to K^+^ restriction

Since muscles participate in the K^+^ balance by releasing their intracellular K^+^ into the extracellular compartment, we first measured the muscle K^+^ contents of the WT and GDF15-KO mice under normal or K^+^ depleted conditions.

As shown in Figure 5A, under normal conditions, muscles (gastrocnemius) of WT mice contained 20% significantly more K^+^ than those from GDF15-KO mice. This result suggests that the GDF15-KO mice already compensate the lack of this factor by releasing muscle K^+^. As expected, the muscle K^+^ content is significantly decreased in WT mice after 4 days of K^+^ restriction compared with the normal condition (66±3 vs 82±2 μmol/g, respectively). Conversely, GDF15-KO mice that have already a low muscle K^+^ content under normal condition are unable to decrease it even more (68±3 vs 61±5 μmol/g). We then investigated whether the muscle structure was modified in response to K^+^ restriction and found that, in this condition, WT muscles have larger fibres than GDF15-KO mice (2148±15 μm^2^ and 1706±11 μm^2^, respectively, Figure 5B). To confirm that K^+^ restriction differently impacts muscle structure in WT and GDF15-KO mice, NMR imaging of hindlimbs have been carried out to precisely measure the size of the muscles. As shown in Figure 5C, the difference of size in the same animals between normal K^+^ condition and 4 days of K^+^ restriction indicated that the size of GDF15-KO mice is 2-times more reduced compared with the WT mice. These two results (laminin labelling and NMR imaging) strongly suggest that GDF15-KO mice lose muscles in response to K^+^ restriction whereas WT mice, that have the ability to release K^+^ from their muscles, maintain their muscle integrity. We, therefore, measured markers of muscle degradation and showed that plasma creatine kinase is more elevated in GDF15-KO mice compared with the WT mice (Figure 5D) after 4 days of K^+^ restriction. In addition, the muscle expression of two genes related to muscle atrophy, TRIM63 and Fxbo32 (27), was found strongly upregulated under low-K^+^ in GDF15-KO mice diet compared with the WT mice (Figure 5E-F).

**Figure 5:**
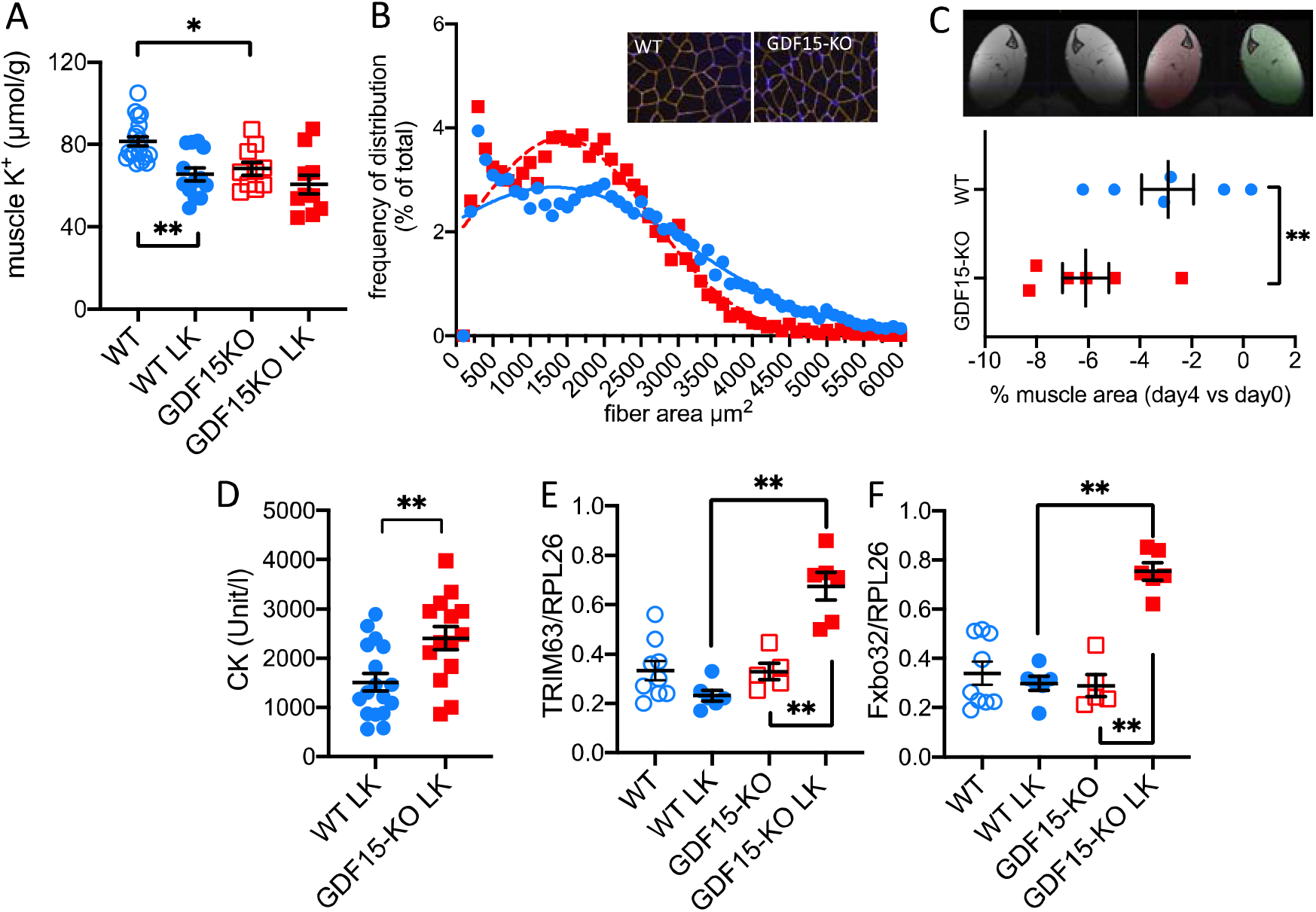
Muscle differentially contributes to K^+^ balance in WT and GDF15-KO mice. **A**/ Muscle K^+^ content from gastronecmius of WT (blue circles) and GDF15-KO (red squares) mice under normal diet (empty symbols) and LK diet (day 4, filled symbols). Results are shown as mean±SEM (n=5) and analyzed by one way ANOVA test and Tukey’s post-test (* p<0.05). **B**/ Distribution of the frequency of muscle fiber area in quadriceps of WT (blue circles, plain line) and GDF15-KO (red squares, dotted line) mice under LK diet for 4 days (bin width 100 μm^2^, n=3 mice, 2 slices/mouse, 8197 and 8734 fibers analyzed in WT and GDF15-KO mice, respectively. Gaussian curves were fitted to the data of WT and GDF15-KO mice, respectively. **C**/ Example of the NMR images of the hindlimb muscles from a WT mouse under low-K diet. Colored portions correspond to the region of interest (ROI) used for calculation of the muscle area. Measurements were performed in WT (circles) and GDF15-KO (squares) mice in normal K^+^ diet and after 4 days of low-K^+^ diets. For the same animals, the differences of the muscle surfaces, between the day before diet modification and the day 4 of the LK diet, were calculated. Results are shown as mean±SEM (n=6) and analyzed by a non-paired Student t-test (*p<0.05). **D**/ Plasma creatine kinase (CK) activity of WT (circles) and GDF15-KO (squares) mice under LK diet (day 4). Results are shown as mean±SEM (n=14) and analyzed by non-paired Student t-test (* p<0.05). **E** and **F**/ mRNA expression of TRIM63 and Fxbo32 in muscles from WT (circles) and GDF15-KO (squares) mice under LK diet (day 4). Results are shown as mean±SEM (n=6) and analyzed by and analyzed by one way ANOVA test and Tukey’s post-test (** p<0.01).

## Discussion

GDF15 is generally expressed at a low level under normal/healthy conditions whereas its plasma concentration or urine excretion increase significantly in front of different stresses that generally involve mitochondria dysfunction (28–30). Among its different roles that have been demonstrated (for review see (14)), its action as an anorectic factor that induces loss of food intake and weight is of major importance (31, 32). Interestingly, aging and cachexia can also induce GDF15 production (33) that may trigger a vicious circle, leading to decrease of appetite which, then, aggravates the situation. GDF15 is also described as a factor that plays a role as negative feedback in response to obesity, limiting energy intake and storage (for review, see (34)). In addition to these global metabolic effects, GDF15 also became a marker of solid cancer tumor progression with the potential to be used as a prognostic factor (for review (35)). Regarding kidney, GDF15 has been recently identified in chronic kidney disease as pro-cachectic factors (36) and treatment of mice with GDF15 peptide following unilateral ureter obstruction decreases renal fibrosis (37).

### GDF15 regulates renal ion excretion

We recently added to this list of functions, the involvement of GDF15 in the control of ionic balances. Indeed, we described that GDF15 expression was strongly increased in kidney collecting ducts in response to acid load (12) and showed that it controls the proliferation rate of AIC in this situation (13). The mechanism of this system was recently elucidated, showing that GDF15 was produced by the principal cells through a vasopressin-dependent pathway and then could activate directly or not ErbB2 receptor on AIC to trigger cell division (25). In the present study, we showed that GDF15 expression by collecting duct is not specific of metabolic acidosis but is also induced by K^+^ restriction and was more systemic, with increased expression in all renal segments and different gut parts, leading to an increase of the urinary and the circulating levels of GDF15. Using samples from a recent study where we tested the adrenal response of healthy volunteers to a mild decrease of K^+^ intake (3), we did not measure a significant increase of GDF15 in the urine of the participants which may be due to the fact that we only moderately decrease the K^+^ intake by a factor 2. However, this challenge was sufficient to show that both urine GDF15 and the plasma K^+^ level of the participants were correlated. Interestingly, in the K^+^ depleted group, the individuals that maintain a normal K^+^ value were those that had elevated urine GDF15 levels, indicating that the ability to increase GDF15 could protect against hypokalemia. Further investigation with an increased number of volunteers would be required to better define the role of GDF15 in human, but this result already confirms the involvement of GDF15 in the K^+^ balance in human too.

Acidosis and K^+^ restriction are well known conditions leading to increase the number of AIC. There are evidences in the literature for an increase of the proliferation rate of these cells in both conditions (10, 13, 25, 38) but also for interconversion from one type of cell (principal or ICB) into ICA (39–41). Here, we cannot definitively conclude for the involvement of GDF15 in one mechanism or another, but we believe that the apparition of doublets of ICA and their reduction in its absence rather plead for a role in the proliferation of ICA. The absence of GDF15 impedes the increase of ICA number in response to K^+^ restriction, by a mechanism that, as in acidosis (25), involves ErbB2 receptor. In most of the peripheral tissues such as kidneys, the GDF15 receptor GFRAL is absent (42), indicating that the peripheral action of this factor must be transduced by another receptor. Whether GDF15 directly binds to ErbB2 receptor remains to be clearly establish, however, it has already been involved in the activation of this receptors in different types of cancer (43–45) where it promotes cell proliferation and pErbB2 was successfully immunoprecipitated with GDF15 (46). The consequences of this inadaptation are a loss of K^+^ in the urine and a rapid decrease of the plasma K^+^ level, indicating that GDF15-mediated AIC proliferation is of major importance in response to K^+^ restriction.

### Extrarenal role of GDF15 in response to K^+^ depletion

The finding that induction of GDF15 is not restricted to a particular renal segment but also to gut parts is in good agreement with the recent observation that after treatment with metformin, an anti-diabetic drug (47) the source of GDF15 was the ileum and the distal colon (48). There are also many examples showing that muscle produces GDF15 in different contexts such as exercise (49) or cachexia (33). In our experiments, after 4 days of dietary K^+^ restriction, the expression of GDF15 by muscles was not significantly increased but the tendency we observed suggest that further experiments should be performed to investigate whether the duration of the diet could influence the response of this tissue.

It is known for years that the gut has the ability to sense K^+^ content and may release factors that permit to rapidly adapt the urine K^+^ excretion without any modification of the plasma K^+^ level. The elegant works, mainly by JH Youn and AA McDonough, using the “K^+^-clamp” technic were recapitulated in (50) and clearly established the concept of feedforward control of the K^+^ balance. A that time, the proposed factors involved in the feedforward control system were not formally identified and were supposed to “inform” the kidney in a situation of K^+^ loading to increase its excretion in the urine. We propose that this system, through GDF15, may also contribute to activate renal K^+^ retention mechanisms to prevent a drop in plasma K^+^ level in case of low K^+^ intake.

Muscles also contribute to maintain a normal plasma K^+^ value by releasing intracellular K^+^ (6, 51) through the regulation of Na,K-ATPase isoforms (52). However, GDF15-KO mice have a low muscle K^+^ content at baseline and seems not able to reduce it even lower. Instead, we observed that GDF15-KO mice decreased their muscle volume through a GDF15-independent mechanisms. A simple explanation is that a low intracellular K^+^ concentration is associated to cell death (53), because it would hyperpolarize the membrane and activates Cl^-^ secretion, cell shrinkage and ultimately apoptosis. Therefore, in GDF15-KO mice, since their level of intracellular K^+^ is already low, the release of K^+^ from muscle cells, as expected in response to K^+^ restriction, rapidly leads to muscle cell shrinkage and destruction. In our study, we cannot distinguish between cell shrinkage and cell death to explain the decrease of muscle volume, but probably both processes are involved since we observed the presence of markers of muscle lysis. The muscle lysis is inefficient to maintain the plasma K^+^ value in a normal range since the kidneys of GDF15-KO mice do not retain it efficiently. Therefore, this is an example where the absence of GDF15 leads to a loss of muscle whereas GDF15 is in, other circumstances, described as a muscle atrophic factor (15, 36).

Altogether, GDF15 appears to be a factor that control plasma K^+^ level by regulating the renal cell plasticity. Its absence inducing a compensatory mechanism that leads to muscle atrophy.

## Material and Methods

### Animals

Experiments were performed on C57BL/6J wild-type and knock-out mice for the GDF15 (GDF15-KO). The GDF15-KO mice were first provided by Dr. Se-Jin Lee (John Hopkins University, Baltimore, MD) and backcrossed with C57BL/6J (13). The animals were kept at CEF (Centre d’Explorations Fonctionnelles of the Cordeliers Research Center, Agreement no. A75-06-12).

### Human urine samples

The samples used to measure urine GDF15 were described previously in (3). Briefly, The healthy volunteers were Caucasian males 18 to 35 years of age. Inclusion criteria were BMI ranging from 18 to 30 kg/m^2^, plasma ionogram and liver panel within normal range, estimated glomerular filtration rate (MDRD) > 60 ml/min/1.73 m^2^. They were depleted in K^+^ by treatment with 30 g daily sodium polystyrene sulfonates (Kayexalate®, Sanofi-Adventis France) over two days, followed by 5 days under a low-potassium diet (25 mmol/d).

### Metabolic analysis

To record physiological parameters, mice were placed in metabolic cages (Techniplast, France) and were fed a standard laboratory diet (0.3 % Na ^+^ and 0.6 % K^+^; UPAE, INRA, Jouy-en-Jossas, France) or a low-K^+^ diet (0.28% Na^+^ and 0.01 % K^+^; UPAE, INRA, Jouy-en-Jossas, France). Mubritinib (Euromedex, Souffelweyersheim, France) treatment was performed by gavage twice daily (8 mg/kg/day) for 4 days along with a normal or a low-K^+^ diet.

Urinary creatinine concentrations and plasma ASAT were determined using an automatic analyzer (Konelab 20i; Thermo, Cergy Pontoise, France). Urinary K^+^ concentration was determined by flame photometry (M420, Sherwood Scientific, France). Plasma K^+^ was measured by retro-orbital puncture on the anesthetized animal (a mix of xylazine, 10mg/kg and ketamine 100mg/kg) with an Epoc pH/blood-gas analyzer (Siemens Healthineers, Saint Denis, France). Plasma and urine GDF15 were measured by ELISA test (MG150 and DGD150, for rodent and human GDF15, respectively, R&D Systems) and tissue GDF15 was measured using an ELISA test (Elabscience).

### Quantitative PCR

After collection, mRNA of kidney, muscle and part of the gut were extracted by the TRI reagent (Invitrogen, Villebon sur Yvette, France) following the manufacturer’s instructions. Isolation of 40-60 renal segments was performed according to localization in the kidney (cortex vs medulla) and well-defined morphologic characteristics under binocular loupes after kidney treatment with Liberase (Sigma-Aldrich, St Quentin Fallavier, France) as described in (25). RNA extraction from these segments was performed using RNeasy micro kit (Qiagen, Hilden, Germany). mRNA was then retro-transcribed into cDNA (Roche Diagnostics, France) according to the manufacturer’s instructions and real-time PCRs were performed on a LightCycler (Roche Diagnostics, France). No signal was detected in samples that did not undergo reverse transcription or in blank runs without cDNA. In each run, a standard curve was obtained using serial dilution of stock cDNA.

### Immunolabelling on isolated OMCD for cell counting

15-20 short OMCD segments were isolated from Liberase-treated kidney and transferred onto a Superfrost Plus glass slides and treated as described recently in (25). AIC were identified as AE1+ cells after labeling with an anti-AE1 antibody (1/500, gift from C.A. Wagner). For cell and doublet counting, a 3D reconstruction from all stacks (ImageJ) was performed from images of AE1-labelled OMCDs acquired by confocal microscopy (25).

### Immunolabelling for laminin on muscles

Quadriceps were collected and fixed in 4% paraformaldehyde (PFA) solution for 1 hour, rinsed in PBS for 30 min and immediately frozen in optimum cutting temperature medium (OCT). Four μm thick sections were cut with a cryostat (Leica CM305S). Before labeling with a rabbit anti-laminine (1/400 Abcam, ab11575) and a secondary anti-rabbit antibody (Alexa Fluor 555, Abcam), the slices underwent a retrieval procedure (Tris-EDTA pH9, 10 min, 95°C). The labeled slices were then observed using an Axio Scan Z1 (Zeiss) microscope allowing acquisition of the whole slice. The muscle fiber areas were determined using the QuPath software (54) and the frequencies of distribution were assessed using GraphPad Prism software.

### NMR imaging of mouse hindlimbs

NMR experiments were performed on WT and GDF15-KO mice (n=6) at baseline and 4 days after Low-K^+^ diet. NMR data were acquired using a 7T Bruker BioSpec system interfaced with an Advance III spectrometer (Bruker BioSpin MRI GmbH, Ettlingen, Germany). Mice were scanned under isoflurane anaesthesia (1-2%, 1l/min O2) on a water-heating pad. Respiratory frequency was maintained between 80-100 motions/min. The NMR setup used was a ^1^H transceiver surface Cryoprobe placed next to the anterior compartment of the two hindlimbs. Muscle atrophy was evaluated from the maximum cross-sectional area of the legs (CSAmax) measured on one slice from an axial high-resolution (50*50*200 μm^3^) gradient-echo images (12 slices, slice gap = 0.5 mm, TE = 3.66 ms, TR = 166.26 ms, FA = 25°, NA = 12, acquisition time = 8 min 18s). Contours of the legs were drawn using ITK-SNAP 3.8.0 (Free Software Foundation, Inc) software.

### Data and statistical analysis

Results are expressed as mean±SEM. The numbers of mice used in these experiments are indicated in the legends (n). Normality of the result distribution is tested using the Shapiro-Wilk analysis. Non-parametric (One-way or two-way ANOVA tests) or parametric (Student t-test) tests were used to determine statistical significance (see the legends), differences with p<0.05 were considered significant.

### Study approvals

All animal experimentations were conducted in accordance with the institutional guidelines and the recommendations for the care and use of laboratory animals put forward by the Directive 2010/63/EU revising Directive 86/609/EEC on the protection of animals used for scientific purposes (project has been approved by a user establishment’s ethics committee and the Project Authorization: number 21927).

Regarding the use of human samples, the study was approved by the local Ethics Committee (P120906 - CPP Ile de France VI, NCT02297048) and informed consent were obtained from the volunteers before participating in the study (registered on clinical trial registry, NCT02297048).

## Author contributions

SL, CW, CR, LM, BM, LC: conducting experiments, acquiring data, analyzing data

AD, NP, AB, YF, GC, LC: designing research studies, analyzing data

SL, LC, GC: writing the manuscript.

## Acknowledgements

Physiological studies have been performed with the help of Gaëlle Brideau and Nadia Frachon from the “plateforme d’exploration fonctionnelle du petit animal” of the team “Physiologie Rénale and Tubulopathies” at the Centre de Recherche des Cordeliers. We are grateful for the technical assistance of the Centre d’Exploration Fonctionelle crews in the management of our colony of mice. This study was supported by the Agence National de la Recherche (ANR) project ANR-21-CE14-0040-01. S.L. is supported by the Ministère de l’Enseignement Supérieur, de la Recherche et de l’Innovation.

